# Lactic acid bacterium *F. sanfranciscensis* impairs fitness of yeast *M. humilis* in a synthetic wheat sourdough medium

**DOI:** 10.64898/2026.04.01.716005

**Authors:** Anna Elizabeth Wittwer, Diego Segond, Thérèse Marlin, Céline Serre, Amanda Li, Delphine Sicard, Kate Howell

**Affiliations:** School of Agriculture, Food, and Ecosystem Sciences, Faculty of Science, The University of Melbourne, Victoria, Australia; SPO, Univ Montpellier, INRAE, Institut Agro, Montpellier, France

**Keywords:** *Maudiozyma humilis*, *Fructilactobacillus sanfranciscensis*, microbial interactions, wheat sourdough, microbial fitness

## Abstract

Sourdough starters contain simple microbial communities typically consisting of a few bacterial species and one or two yeast species. The yeast *Maudiozyma humilis* and the lactic acid bacterium *Fructilactobacillus sanfranciscensis* often co-occur in sourdough starters and have been presumed to exist in a trophic relationship supported by glucose cross-feeding. However, previous research has highlighted a lack of evidence showing that yeast strains consume the glucose that *F. sanfranciscensis* produces. We have investigated the interaction between sourdough isolates of *M. humilis* and *F. sanfranciscensis* in a synthetic wheat sourdough medium, allowing us to control substrate composition and use flow cytometry to enumerate living and dead cells. *M. humilis* fitness was found to be lower in co-culture with *F. sanfranciscensis* than when grown alone. Analysis of spent medium composition highlighted the reliance of *M. humilis* on glucose rather than maltose for growth. Comparisons of predicted and measured co-culture metabolite content also revealed that *F. sanfranciscensis* consumed less maltose in co-culture than when grown alone. For the first time, we examined potential amino acid cross-feeding between *M. humilis* and *F. sanfranciscensis*, and found that within the pairing, *F. sanfranciscensis* was the main producer of amino acids. Our findings suggest that the *M. humilis*-*F. sanfranciscensis* interaction is likely to be neutral, or even competitive, with the strain identity of *F. sanfranciscensis* playing a defining role in the observed dominance of the bacteria and spent medium metabolite composition.

**Importance:** The association of the yeast *Maudiozyma humilis* and the bacterium *Fructilactobacillus sanfranciscensis* in sourdough starters is well-documented, and together this pairing makes key functional and organoleptic contributions to the final bread product. Their relationship has historically been thought to be stabilised by cross-feeding of glucose to *M. humilis*. However, this theory has been drawn into question by recent research which found no evidence that *M. humilis* consumes the glucose produced by *F. sanfranciscensis*. Our understanding of cooperation, coexistence, and competition in microbial consortia affects approaches to ecosystem management in a broad variety of applied fields. The significance of our research is in demonstrating that this pairing does not interact mutualistically within a specified setting, providing support for neutral or competitive interactions as drivers of ecological stability.

## Introduction

Food ecosystems are one of many environments in which fungi and bacteria co-exist (1). In fermented foods, this combined existence is highly desirable, as the metabolic activity of yeasts and bacteria leads to the accumulation of organic acids, alcohol, CO_2_, and various flavour and aroma compounds (2). Fundamental knowledge of the metabolic interactions of bacteria and yeast is needed to better understand their co-occurrence and interaction stability over time. Moreover, a deeper understanding of how one fermentation microbe’s inputs and outputs are influenced by another’s can be exploited to develop novel defined starter cultures and modulated foods with improved organoleptic and/or health properties.

Yeasts and bacteria always co-occur in type I sourdough starters used to make sourdough bread. Also known as traditional sourdough starters, these are mixtures of flour and water that ferment ‘spontaneously’ (i.e., without inoculum), and over time and with repeated refreshments of the substrate, they become home to stable microbial communities. The typical sourdough starter ecosystem is relatively simple, containing one or two yeast and bacteria species, and not containing more than 6 species in total (3). These micro-organisms carry out the necessary bread-making functions of leavening and flavour development, while certain species offer particular technological and health benefits (4). *Maudiozyma humilis* (formerly *Kazachstania* or *Candida humilis*) is the second most common yeast species found in sourdough starters after *Saccharomyces cerevisiae* (5, 6), and research into its properties has grown in recent years due to an increasing interest in non-conventional yeasts. Importantly, it displays a well-documented co-occurrence with the lactic acid bacterium *Fructilactobacillus sanfranciscensis* in type I sourdough starters (3, 5–10).

There are several possible explanations for this co-existence. Firstly, *M. humilis* is a maltose-negative yeast, despite maltose being the most readily available small sugar in bread dough (11). *F. sanfranciscensis*, however, constitutively expresses a maltose phosphorylase, and is able to metabolise maltose into glucose and glucose-1-phosphate.The glucose resulting from this would ostensibly be available for yeast to consume, thereby benefiting *M. humilis* (12). This cross-feeding hypothesis is typically supported by the widespread co-occurrence of both species in sourdough starters as well as their complementary carbohydrate metabolisms (5, 13, 14). However, experimental data showing that *M. humilis* consumes the glucose produced by *F. sanfranciscensis* is sparse. Additionally, *Saccharomyces cerevisiae* and LAB have been shown to exist in a mutualistic relationship due to amino acid overflow that facilitates the survival of the LAB (15). *Maudiozyma exigua* also only partially competes with LAB for amino acids and facilitates LAB growth via amino acid secretion (16). Due to the phylogenetic similarity *M. exigua* and *M. humilis* (and *S. cerevisiae* to a lesser extent), it is thought that *M. humilis* may be similarly capable of supporting LAB growth through the provision of amino acids.

While the major theory explaining the co-existence and presumed interaction between *M. humilis* and *F. sanfranciscensis* is glucose cross-feeding, heightened levels of glucose have been found in doughs fermented solely by obligately heterofermentative lactic acid bacteria (LAB) and those co-fermented by *M. humilis* and obligately heterofermentative LAB (17). This led authors to conclude that “…there was no clear evidence that yeast strains consume the glucose produced by the obligately heterofermentative LAB” (17). Furthermore, all yeast strains tested exhibited significantly reduced fitness when co-cultured with LAB strains compared to monoculture levels, whereas the fitness of the LAB strains was seldom affected by the presence of a co-culture yeast (17). In a similar experiment, *M. humilis* was shown to neither worsen nor improve *F. sanfranciscensis* growth (18).

To date, a small number of studies have modelled growth of *F. sanfranciscensis* and *M. humilis* co-cultures in simulated sourdough fermentation conditions (19, 20) and in bread dough (10, 17). The present study aims to expand on these studies by testing co-culture performance in a wheat synthetic sourdough medium (WSSM) rather than dough. This enables the use of flow cytometry to quantify both viable and dead yeast and bacterial cells, as well as to quantify the carbohydrates and amino acids present. As no previous studies have examined the possibility of amino acid cross-feeding between *M. humilis* and *F. sanfranciscensis*, a key objective of the present study is to do so alongside carbon metabolite exchange patterns. In doing so we aim to measure how co-cultivation affects the growth performance and thus fitness measured as number of viable cells of *M. humilis* and *F. sanfranciscensis*.

## Materials & Methods

### Strains and pre-culture preparation

Strains used in this study were previously isolated from sourdoughs. Eight previously identified *M. humilis* strains and eight *F. sanfranciscensis* strains were used (**Table 1**). Four strains each of *M. humilis* and *F. sanfranciscensis* were sourced from France and Australia. The French and Australian collections were each composed of four yeast-bacteria pairs, and each of those pairs came from sourdough provided by the same bakery. This was done to minimise potential effects arising from strain variation between sourdough starters as much as possible, and to account for potential selection for cross-feeding or competitive avoidance as a result of the strains having co-evolved (21). However, the Australian collection had only two complete single-sourdough pairs, so two were composed of *M. humilis* and *F. sanfranciscensis* sourced from different sourdoughs. A bakery-derived *S. cerevisiae* strain was used as a reference strain in one experiment. Each strain code indicates its country of origin, the bakery it came from, and a number to differentiate it from other strains from the same origin.

**Table 1.** Species, identifying codes, and origins of yeast and bacterial strains used in this study. *M. humilis* population genomic analyses conducted by Lebleux *et al*. (22).

| Type | Species | Code | Bakery | Region of origin | *M. humilis* genetic diversity (clade no.) |
| --- | --- | --- | --- | --- | --- |
| Yeast | *Maudiozyma humilis* | Aus_LbrY1 | Loafer Bread (rye starter) | Fitzroy North, VIC, Australia | 1 |
|  |  | Aus_NrkY1 | Kate Howell | Natimuk, VIC, Australia | 5 |
|  |  | Aus_OsrY1 | Ovens St. (rye starter) | Brunswick, VIC, Australia | 5 |
|  |  | Aus_OsY1 | Ovens St. | Brunswick, VIC, Australia | 5 |
|  |  | Fr_B10.20F | B10 | Forcalquier, Alpes-de-Haute-Provence, France | 6 |
|  |  | Fr_B17.25 | B17 | Liomer, Hauts-de-France, France | 6 |
|  |  | Fr_B5.TP1.20 | B5 | Noailles, Hauts-de-France, France | 6 |
|  |  | Fr_B6.15 | B6 | Barbèrey-Saint-Sulpice, Grand Est, France | 4 |
| Bacteria | *Fructilactobacillus sanfranciscensis* | Aus_KbB1 | Ket Baker | Wallington, VIC, Australia |  |
|  |  | Aus_OsB1 | Ovens St. | Brunswick, VIC, Australia |  |
|  |  | Aus_OsRB1 | Ovens St. (rye starter) | Brunswick, VIC, Australia |  |
|  |  | Aus_RB2a | Red Beard Bakery | Trentham, VIC, Australia |  |
|  |  | Fr_B10.CP109 | B10 | Forcalquier, Alpes-de-Haute-Provence, France |  |
|  |  | Fr_B17.CIRM2 059 | B17 | Liomer, Hauts-de-France, France |  |
|  |  | Fr_B5.LB.102 | B5 | Noailles, Hauts-de-France, France |  |
|  |  | Fr_B6.AG.102 | B6 | Barbèrey-Saint-Sulpice, Grand Est, France |  |
| Yeast | *Saccharomyces cerevisiae* | Fr_MTF 3622 |  |  |  |

*F. sanfranciscensis* strains were maintained at -70 °C in de Man-Rogosa-Sharpe (MRS) or MRS-5 medium (23) with 15% v/v glycerol, and *M. humilis* strains and *S. cerevisiae* were maintained in yeast-peptone-dextrose (YPD) medium with the same concentration of glycerol.

Prior to use in experiments, bacteria were pre-cultured in MRS-5 broth in anaerobic conditions at 25 °C for 24 h with no agitation, and yeasts were grown in YPD broth under constant agitation (220 rpm) at 28 °C for the same amount of time (23).

### Wheat synthetic sourdough medium (WSSM)

WSSM was prepared as described in Boudaoud *et al.* (24). It contains, per L: 24 g wheat peptone, 4 g KH_2_PO_4_, 4 g K_2_HPO_4_, 0.2 g MgSO_4_.7H_2_O, 0.05 g MnSO_4_.H_2_O, 1 mL Tween 80, and 0.899 L MilliQ water. Prior to autoclaving, the pH was adjusted to 4.5 with 3 M citric acid. Once autoclaved, 100 mL of a filter-sterilised (0.22 μm) sugar solution was added. This mixture was composed of 150 g/L glucose and 350 g/L maltose to achieve a final concentration of 15 g/L glucose and 35 g/L maltose. The medium was completed by the addition of 1 mL filter-sterilised (0.22 μm) vitamin solution containing 200 mg/L each of cobalamin, nicotinic acid, folic acid, pantothenic acid, pyridoxal-phosphate, and thiamine.

Modified forms of the WSSM were also prepared to test the effects of pH and sugar composition on growth, measured as optical density. WSSM-G containing only glucose and WSSM-M containing only maltose were also prepared as above but with a final concentration of 50 g/L glucose or maltose. WSSM adjusted to pH 3, 4, 5, and 6 ± 0.05 with 20% lactic acid was also prepared to test the range of pHs commonly observed in dough during fermentation (25).

### Mono-culture growth curves

Growth kinetics of all mono-cultures were investigated using growth curves.

The effect of varied carbohydrate composition on the growth of all strains (**Table 1**) was assessed using WSSM, WSSM-G and WSSM-M. Standardised broth cultures were prepared as follows. Precultures were divided into three, centrifuged at 4500 rpm for 5 min, and the preculture medium removed and replaced with one of the three WSSM types to replace the volume of growth media. Aliquots of the resulting cell suspension were diluted 9:1 with distilled water and the optical density at 660nm was measured against test medium in distilled water as the blank. The quantity of medium required to standardise each preculture to OD_660_ = 0.1 was calculated and added. Aliquots of 200 μL standardised precultures were added to a 96-well flat-bottomed plate in triplicate and uninoculated medium was added to serve as blanks. Optical density at 660nm was measured in a plate reader (CLARIOstar® Plus, BMG LABTECH) every 30 min for 24 h at 25 °C, with a brief agitation step before each measurement.

One *M. humilis* strain (Aus_OsrY1) was assessed for its growth performance in pH-adjusted WSSM. Pre-culture OD_660_ was standardised to 0.1 in 0.1% peptone water, and 10μL of this standardised cell suspension was added in triplicate to 150 μL of each of the four pH-adjusted WSSM. The OD was measured with the xMark Microplate Spectrophotometer (Bio-Rad) at 660 nm every 30 min for 24 h with shaking for 5 s on low orbital mode right before measurement.

### Co-culture growth assay

To test the effect of co-cultivation on strain fitness and medium composition, each *M. humilis* and *F. sanfranciscensis* strain was cultivated alone (8 mono-cultures) and in a pair with each corresponding yeast/bacterial strain (64 co-culture combinations) in standard WSSM. Blanks consisted of un-inoculated WSSM.

Pre-culture cell counts were performed using an Attune™ NxT Acoustic Focusing Cytometer (Thermo Fisher Scientific). The pre-culture cell density was then standardised to 10^7^ cells/mL for *M. humilis* strains and 10^9^ for *F. sanfranciscensis* strains such that the starting concentration once the WSSM had been added were 10^6^ and 10^8^ cells/mL for the yeast and bacterial strains respectively. This was done to reflect the typical ratio of bacterial cells to yeast cells in a traditional sourdough starter (26). Aliquots of 100 μL of the standardised yeast and/or bacterial precultures were added to deep well V-bottomed plates, and WSSM was added to make up the volume to 1 mL. Plates were incubated at 25 °C for 24 h.

### Flow cytometric assessment of viability of LAB and yeast

After incubation, cultures were mixed by pipetting and 50 μL cell suspension was diluted in 150 μL sterile physiological saline (0.9% w/v NaCl). Saline was used instead of phosphate buffered saline to reduce background noise, allowing the small *F. sanfranciscensis* cells to be detected. Cells suspended in saline solution were centrifuged for 3 min at 3000 rpm. The WSSM was removed (175 μL) and replaced with the same quantity of saline solution. This suspension was diluted 1:200, then stained with 250 nM SYTO 9™ to stain living cells and 1.5 µM propidium iodide (PI) to stain the dead cells. Suspensions were incubated for 15 min at room temperature in the dark.

Flow cytometric analyses were performed with an Attune™ NxT Flow Cytometer equipped with a 488 nm blue laser and a 561 nm yellow laser. SYTO 9™ fluorescence was collected in the BL1 channel (530/30 nm), and PI fluorescence was collected in the YL2 channel (620/15 nm). The flow rate was set to 200 μL/min and acquisition volume was set to 70 μL per sample. Instrument performance was verified using Attune™ Performance Tracking Beads. Instrument settings were optimised using unstained cells and single-stained controls (SYTO 9™ only and PI only). These controls were used to generate a compensation matrix with the Attune™ NxT software by minimising fluorescence spillover between BL1 and YL2 channels. There was no spectral overlap between SYTO 9™ and PI fluorescence in our conditions, however, so the compensation matrix was not applied to the sample analysis.

### Gating strategy

Collected fluorescence data were visualised and gated in Attune™ Cytometric Software 5.3.2415.0 © 2021 (Thermo Fisher Scientific Inc.). *M. humilis* cells were identified using forward- and side-scatter characteristics (FSC and SSC respectively). Because *F. sanfranciscensis* events were mixed with a substantial amount of low-scatter electronic background and debris, a secondary fluorescence gate was applied to exclude non-cellular events. This gate was established using unstained and single-stained controls and included all events exhibiting fluorescence in either the SYTO 9™ (BL1) or PI (YL2) detection channels. This ensured that viable, membrane-compromised and non-viable bacterial cell events were retained while excluding non-fluorescent events. Singlet discrimination was performed using SSC-A versus SSC-H plots to remove doublets and aggregates.

The size and light-scattering properties of *M. humilis* and *F.sanfranciscensis* differed markedly, so cell viability was assessed separately for the yeast and bacterial populations using bivariate SYTO 9™ versus PI fluorescence plots. Quadrant boundaries were established independently for each population using unstained, single-stained, live-cell, and heat-killed controls acquired under identical instrument settings. The following populations were defined: non-viable cells with compromised membrane integrity (Q1, −SYTO 9™/+PI), membrane-damaged or injured cells (Q2, +SYTO 9™/+PI), background events and residual debris (Q3, −SYTO 9™/−PI), and viable cells with intact membranes (Q4, +SYTO 9™/−PI). The gating strategy is summarised in supplementary **Figure S1**.

The assay was then validated with mixtures containing defined proportions of live and dead cells (0, 25, 50, 75, and 100% dead cells).

### Metabolite analysis

Cell suspensions in WSSM were centrifuged at 4500 rpm for 3 min to remove remaining cells, and the supernatant was extracted for metabolite analysis. The un-inoculated blank treatments were also measured to act as a baseline. Concentrations of maltose, glucose, raffinose, glycerol, ethanol, mannitol, pyruvate, acetate, lactate, and 23 amino acids in the spent WSSM were determined by high performance liquid chromatography (HPLC). For carbohydrate and derived compound analysis, the supernatant was diluted with 10 mM H_2_SO_4_ prior to chromatographic analysis (27). Quantification was made with a Rezex ROA column (Phenomenex) set at 60°C on an HPLC 1260 Infinity, Agilent Technologies. The separation was resolved isocratically with 0.0025 mol/L H2SO4 at a flow rate of 0.6 mL/min. Concentrations of acetate, malate, lactate and α-ketoglutarate were measured with a UV detector at 210 nm and other compounds with a refractive index detector. Amino acids were quantified using a Shimadzu Nexera Series system using a C18 Shim-pack XR-ODS II column (2.2 μm, 100 x 3.0 mm). A pre-column derivatisation of amino acids with OPA/MPA and FMOC was performed to then be detected by fluorescence and/or UV (fluorescence detection: OPA complex Ex350nm, Em450nm, FMOC complex Ex266nm, Em305nm and UV detection, OPA complex 338nm, FMOC complex 262nm). Two internal standards (norvaline and sarcosine at 50µM) were added to the standards and samples, which were then filtered through Millipore syringe filters (Millex-GV 0.22 μm PVDF). Metabolite amounts were expressed as g or mg/L WSSM. Three of the amino acids (methionine sulfone, cysteic acid, and hydroxyproline) were not detected in any samples and so were omitted from further analysis.

Medium pH was also measured at 24 h for the inoculated treatments and the un-inoculated blanks.

### Testing for co-culture interaction effects

To look for metabolic interactions between yeasts and bacteria, we used their mono-culture metabolite concentration data to calculate a predicted co-culture value for each metabolite. Here, our calculation assumed that no interactions between the microbes occurred (concentration of metabolite by yeast C_Y_; by bacteria C_B_; **Table 2**). We then compared the predicted (C_CC_) and experimental values (C_Y_; C_B_) for each metabolite to look for differences. We then classified these possible interactions with the scenarios described in **Table 2**.

**Table 2.**
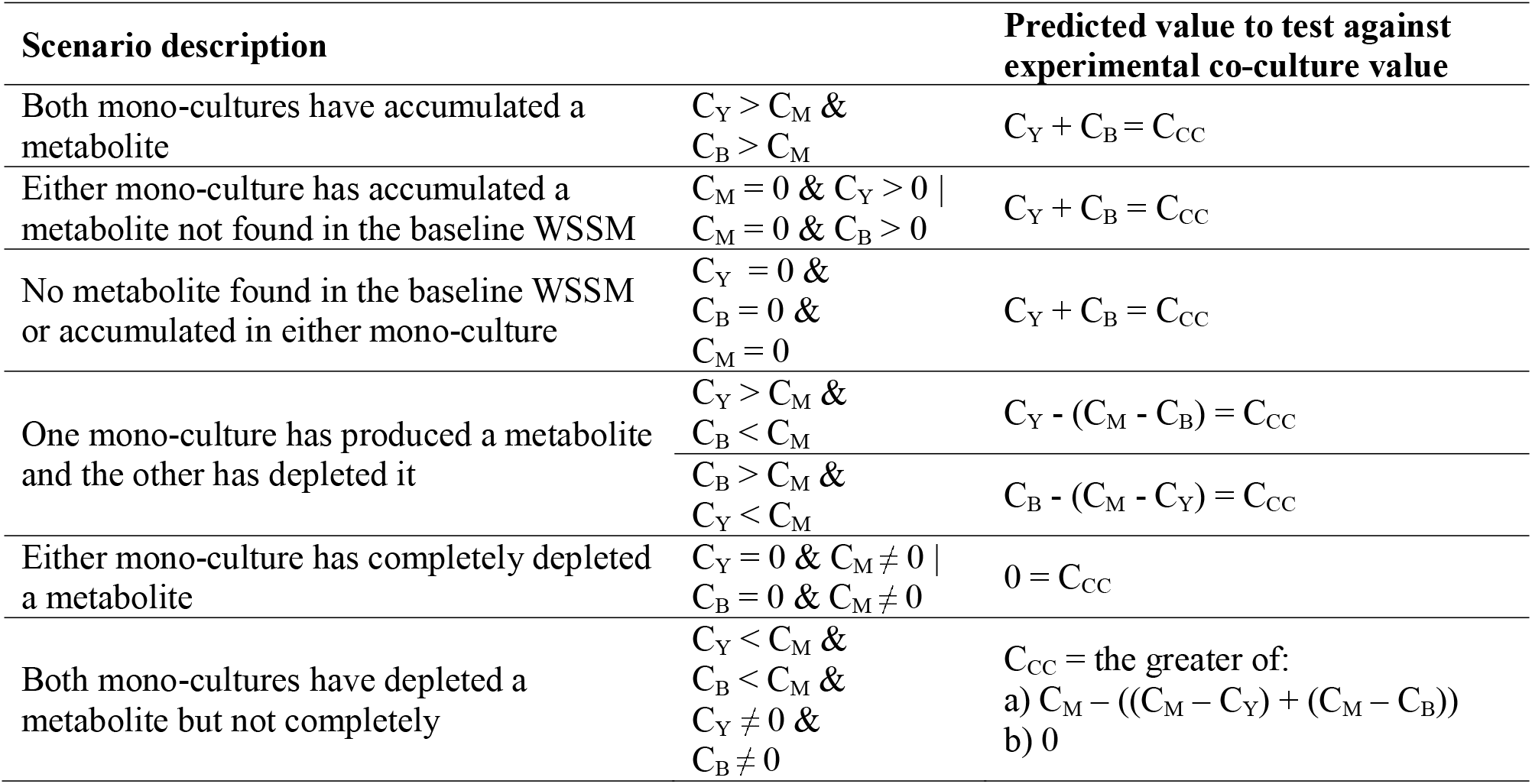
Scenario testing for co-culture interactions using metabolite concentrations in yeast and bacterial monoculture in WSSM. C_M_ = baseline medium (WSSM) metabolite concentration, C_Y_ = yeast mono-culture metabolite concentration, C_B_ = bacterial mono-culture concentration, C_CC_ = predicted co-culture concentration.

### Statistical analyses

Experiments were conducted in triplicate. Group means were compared with t-tests or one-way ANOVA, and pairwise comparisons computed with Tukey’s HSD test at a significance level of α = 0.05. All statistical analyses were performed in R (28).

A schematic summary of the methodology of this study is given in supplementary **Figure S2**.

## Results

The WSSM was inoculated with all *M. humilis* and *F. sanfranciscensis* strains in mono-culture and in all 64 possible pairwise combinations. After 24 h of growth at 25 °C cells were stained and counted by flow cytometry.

Counts of live *F. sanfranciscensis* cells were the highest in both monoculture and co-culture conditions (**Figure 1**A). Counts of live and dead *F. sanfranciscensis* cells did not differ significantly depending on whether they were co-cultivated with *M. humilis*. However, we observed significantly fewer live *M. humilis* cells and more dead yeast cells (data not shown) in co-culture treatments compared to yeast mono-cultures (**Figure 1**B).

**Figure 1.**
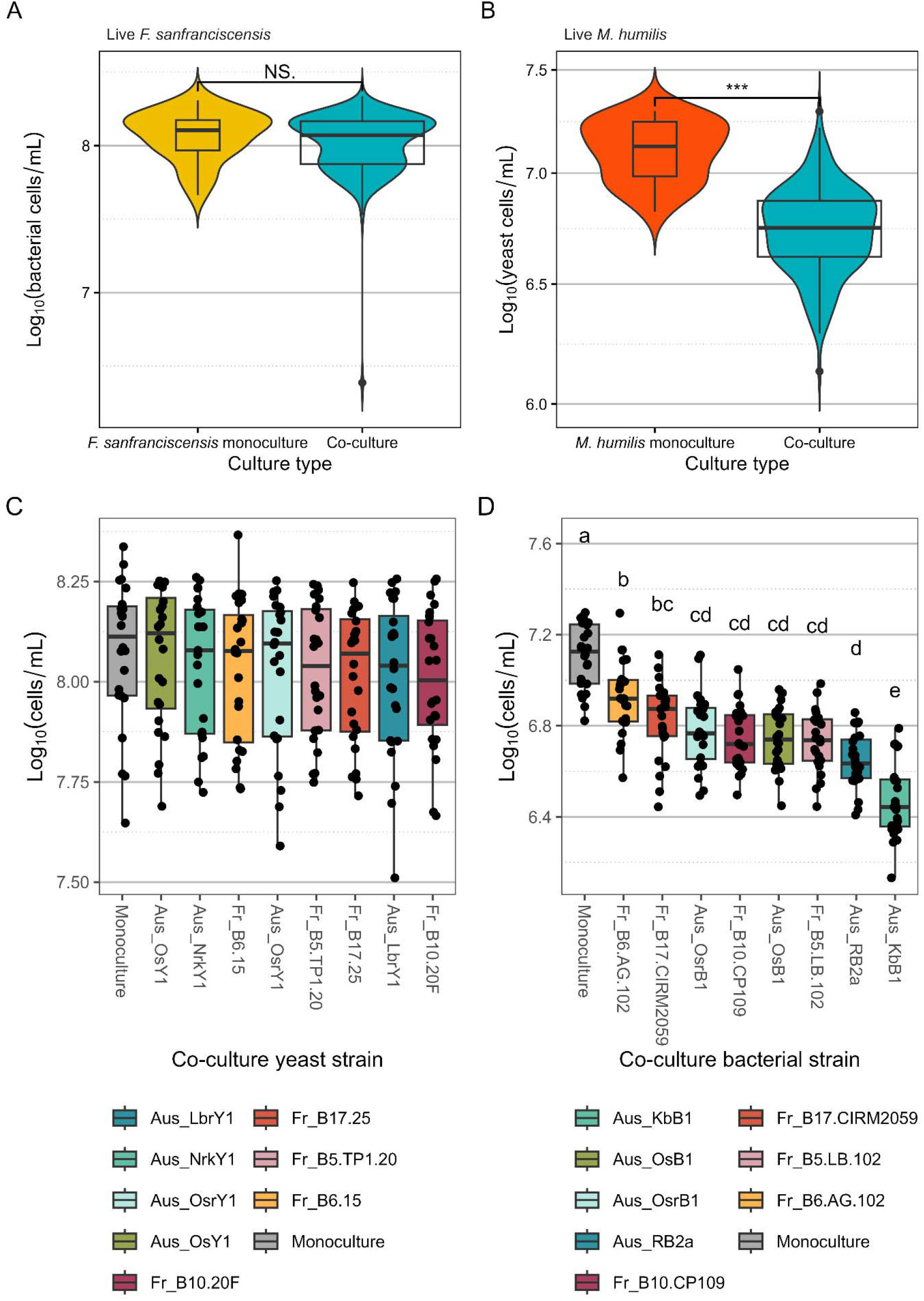
Effect of co-cultivation on fitness of *F. sanfranciscensis* (A) and *M. humilis* (B) overall and according to its accompanying yeast (C) and bacterial (D) strain. Cells in wheat sourdough simulation medium (WSSM) after 24 hours of growth counted by flow cytometry. *** *p* < 0.001, means not sharing letters are significantly different from one another.

We also looked for a strain-based effect on the fitness of *F. sanfranciscensis* and *M. humilis* in co-culture. Counts of live yeast cells varied significantly according to which accompanying *F. sanfranciscensis* strain was present. *F. sanfranciscensis* fitness was not significantly affected by either the presence or strain identity of *M. humilis* in co-culture (**Figure 1**C), but bacterial strains varied significantly in their ability to impair yeast fitness (**Figure 1**D). Live *M. humilis* cell counts were approximately one log-fold lower when grown with *F. sanfranciscensis* Aus_KbB1 than when grown alone.

To investigate the metabolites produced and consumed by our strain library, we collected the spent WSSM after 24 h of growth of the *F. sanfranciscensis* or *M. humilis* monocultures and measured carbohydrate and amino acid concentrations. We then compared them to baseline WSSM levels to show the effect of yeast and bacterial metabolism on medium composition (**Figure 2**).

**Figure 2.**
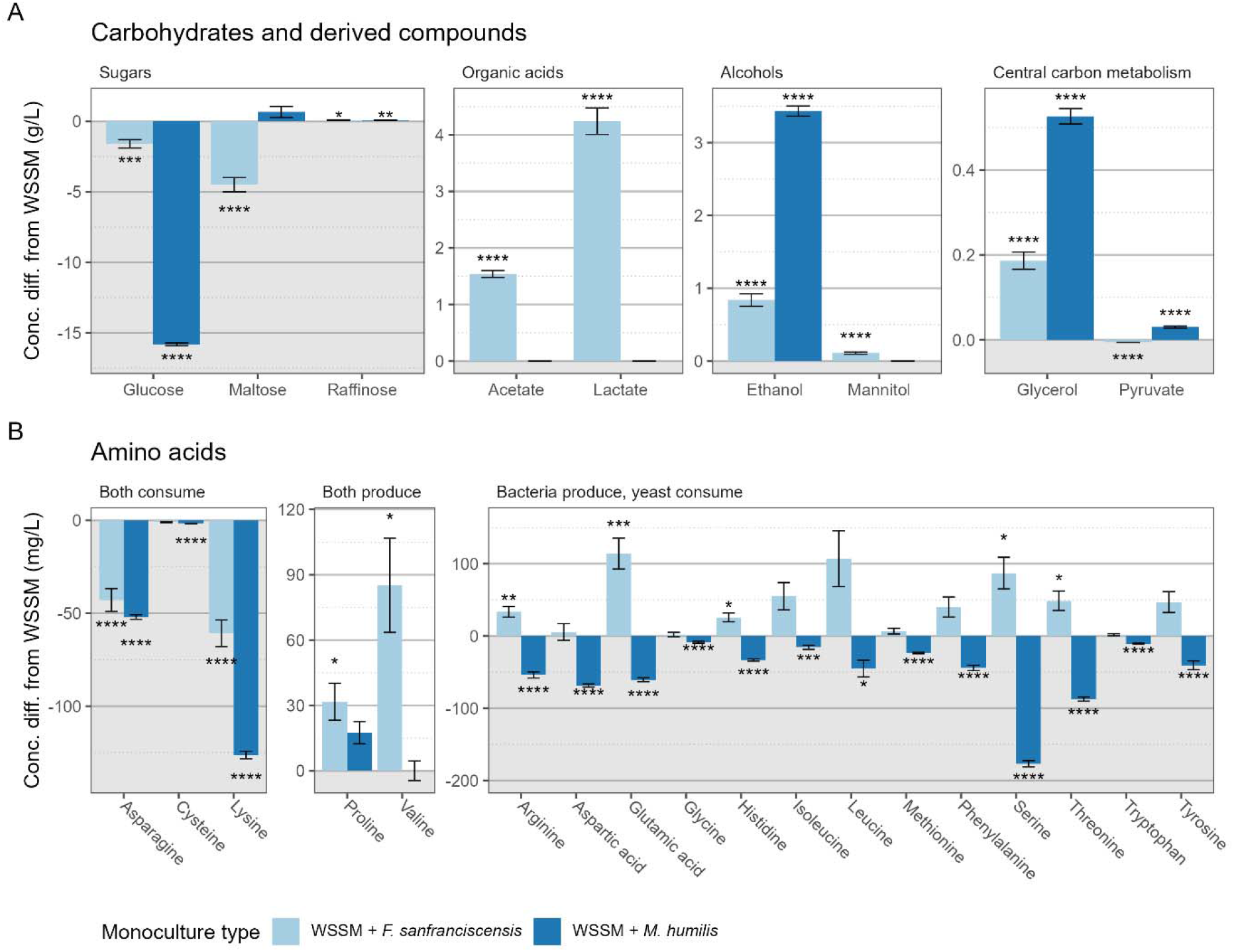
Changes in amino acid (A) and carbohydrate (B) concentrations in wheat sourdough simulation medium (WSSM) after 24 h of growth of *F. sanfranciscensis* or *M. humilis*. * *p* < 0.05, ** *p* < 0.01, *** *p* < 0.001, **** *p* < 0.0001. Statistical comparison of baseline WSSM composition and 24 h monoculture composition by paired t-test with Bonferroni *post-hoc* correction. Error bars represent SEM. Alanine and glycine not shown, no significant differences found.

Glucose was consumed by both *M. humilis* and *F. sanfranciscensis*, but the yeast strains consumed more (**Figure 2**A). Furthermore, while *F. sanfranciscensis* also consumed a small proportion of the available maltose, *M. humilis* consumed none, indicating a reliance of *M. humilis* on glucose in this context. Interestingly, both yeasts and bacteria produced small quantities of raffinose. Only *F. sanfranciscensis* monocultures accumulated organic acids, and lactate production was more than double that of acetate. Both yeasts and bacteria produced ethanol, but yeast produced much more, whereas only *F. sanfranciscensis* produced a small amount of mannitol. Both produced small amounts of glycerol, and only *M. humilis* produced any pyruvate.

Amino acid metabolism patterns fell into three categories: consumption of an amino acid by both *F. sanfranciscensis* and *M. humilis*, accumulation of an amino acid by both, and the production of an amino acid by *F. sanfranciscensis* with the consumption of an amino acid by *M. humilis* (**Figure 2**B). Asparagine, cysteine, and lysine were depleted in both *M. humilis* and *F. sanfranciscensis* monocultures, so we hypothesised that in co-culture, there would be competition for these amino acids. Based on previous findings (15) we also expected that *M. humilis* would accumulate amino acids, and *F. sanfranciscensis* would deplete them. Conversely, we found that arginine, glutamic acid, histidine, serine, and threonine were all significantly higher in the bacterial monoculture than the WSSM, and significantly lower in the yeast monoculture. These amino acids could be involved in a trophic relationship between the two when grown in co-culture (further analysis in **Figure 3**). Yeasts also consumed aspartic acid, glycine, isoleucine, leucine, methionine, phenylalanine, tryptophan, and tyrosine, while increases of the same amino acids in the bacterial monocultures were not statistically significant (**Figure 2**B). In both yeast and bacterial monocultures, proline (17.5 and 31.7 mg/L respectively) and valine (0.053 and 85.2 mg/L respectively) were accumulated, but this was only significant for the *F. sanfranciscensis* monocultures.

**Figure 3.**
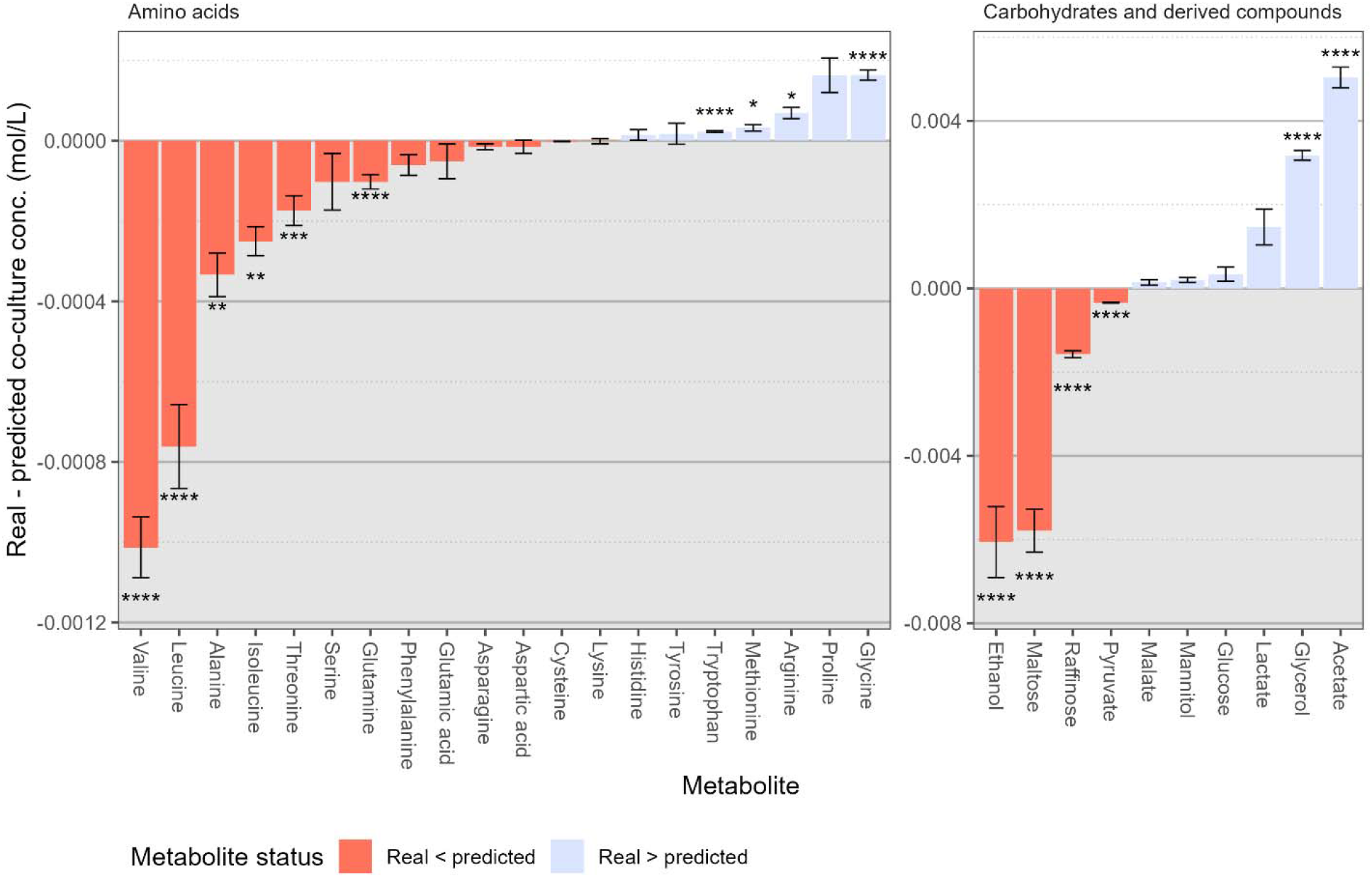
Differences between predicted and real co-culture amino acid and carbohydrate concentrations. * *p* < 0.05, ** *p* < 0.01, *** *p* < 0.001, **** *p* < 0.0001. Statistical comparison of predicted and simulated values by *t*-test with Bonferroni *post-hoc* correction. Error bars represent SEM.

Next, we used these monoculture metabolite concentrations to predict co-culture values (**Table 2**). These predictions were made on the assumption of no interactions occurring between *M. humilis* and *F. sanfranciscensis*, and we compared them to real co-culture metabolite concentrations to search for potential interaction-driven differences (**Figure 3**).

There were several amino acids and carbohydrates whose real co-culture concentrations differed significantly from predicted values (**Figure 3**). Although competition for asparagine, cysteine, and lysine was predicted, no significant differences in concentration were detected between real and predicted values, suggesting that yeast and bacterial consumption of all three amino acids proceeded apace without interference. In real co-culture conditions, amino acid consumption patterns were significantly different from our predictions. Real co-culture threonine concentrations were lower than predicted co-culture values, whereas arginine levels were higher.

This suggests that co-culture conditions increased yeast threonine consumption or reduced bacterial threonine production, and did the inverse for arginine – decreased yeast arginine consumption or increased bacterial arginine production. Yeast consumption of leucine and isoleucine was greater in co-culture than predicted, while yeast consumption of glycine, methionine, and tryptophan was reduced. Although both *M. humilis* and *F. sanfranciscensis* monocultures saw an accumulation of proline and valine (**Figure 2**B), real valine concentrations were significantly lower than predicted.

While no significant differences were found between real and predicted glucose levels, maltose depletion by *F. sanfranciscensis* was increased in real co-culture conditions relative to our predictions (**Figure 3**). Ethanol production was lower than expected in co-culture, whereas glycerol production was increased. Raffinose and pyruvate production were also significantly lower than expected. Bacterial acetate production was significantly higher in real co-culture conditions than our predicted values.

In addition to our examination of strain effects on *F. sanfranciscensis* and *M. humilis* fitness in **Figure 1**C-D, we investigated bacterial strain effects on our fitness, metabolite concentration and pH data (**Figure 4**).

**Figure 4.**
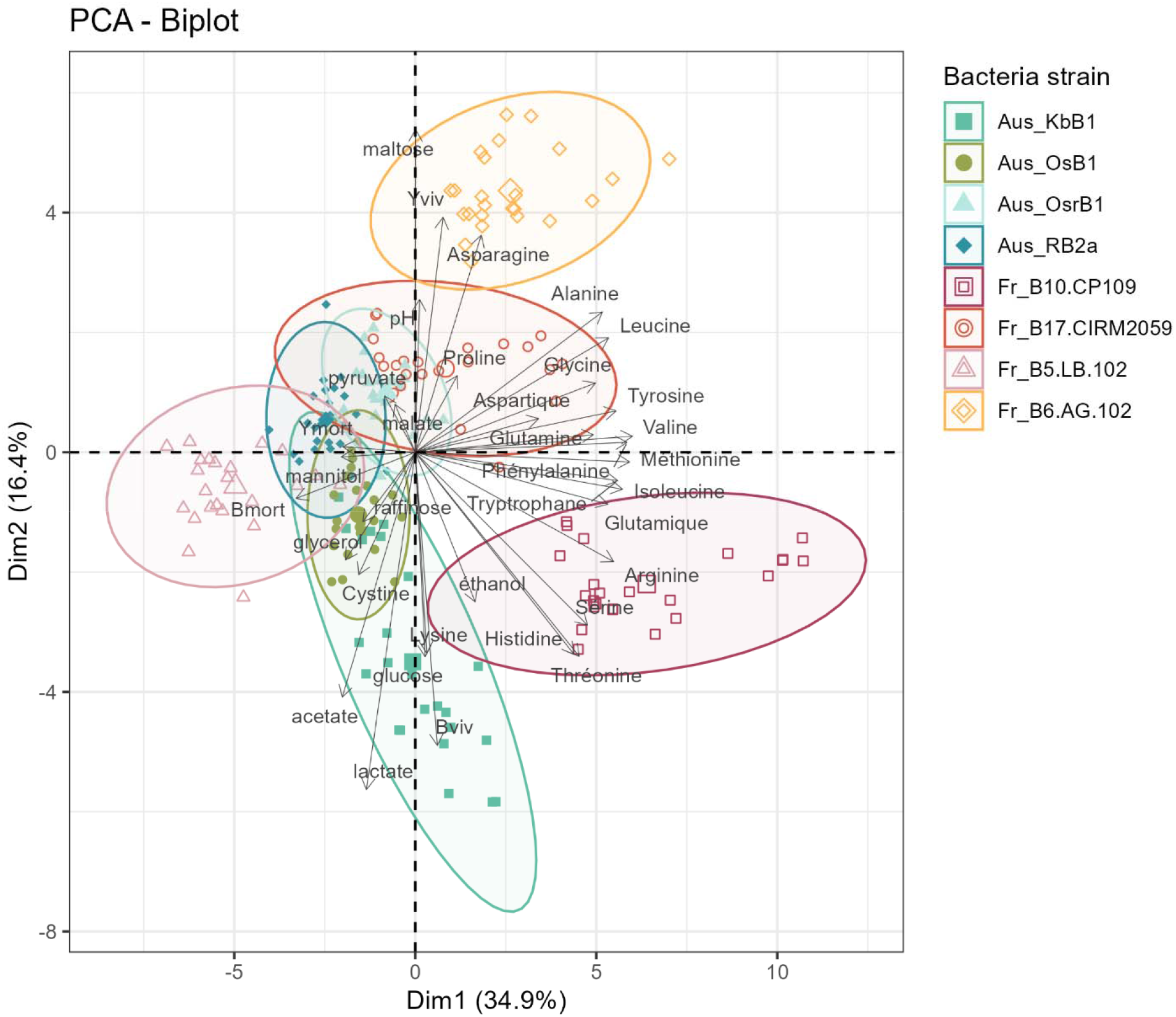
Principal component analysis (PCA) of fitness, metabolite concentration, and pH variables from *F. sanfranciscensis* and *M. humilis* co-culture data. Variables scaled to have unit variance prior to analysis. Yviv: live yeast cell count, Ymort: dead yeast cell count, Bviv: live bacterial cell count, Bmort: dead bacterial cell count.

Principal component analysis showed that 51.3% of the variance between co-cultures could be explained by the first two principal components (**Figure 4**). Axis 1 largely discriminates between Fr_B10.CP109 and Fr_B5.LB.102, and this is mainly linked to amino acids (**Figure 4**), which make the greatest contributions to the principal components. These amino acids are both

Firstly, we wanted to understand whether changes in the pH of the WSSM caused by *F. sanfranciscensis* producing acetate and lactate were impairing *M. humilis* fitness (**Figure 5**A). We found that in WSSM with its pH adjusted to 3, 4, 5, and 6 with lactic acid, *M. humilis* Aus_OsrY1 fitness was not affected. However, we demonstrated that all *M. humilis* strains in our library were negatively affected by the absence of glucose in WSSM (**Figure 5**B), highlighting their dependence on access to usable carbohydrates, especially in comparison with *S. cerevisiae* Fr_MTF 3622. Because data showing that our bacteria were typically producers of amino acids was contradictory to previous findings (15), we wanted to determine whether this phenomenon was being driven by *F. sanfranciscensis* cells dying and autolysing, thereby releasing their contents into the WSSM (**Figure 5**C). We found only a minor correlation between counts of dead *F. sanfranciscensis* cells and total amino acid content (R^2^ = 0.10). Lastly, to investigate the factors underpinning *F. sanfranciscensis* fitness, we examined the relationship between quantities of maltose consumed and lactate produced, as well as associations with counts of live *F. sanfranciscensis* cells and WSSM pH (**Figure 5**D). We found that maltose consumption correlates with lactate production (R^2^ = 0.56), suggesting that maltose uptake is linked by metabolic conversion to lactate output by *F. sanfranciscensis*. This is supported by the absence of maltose consumption and lactate production by *M. humilis* (**Figure 2**A). Furthermore, as maltose consumption and lactate production increase, so too do numbers of living bacterial cells, and this is also loosely tied to WSSM pH.

**Figure 5.**
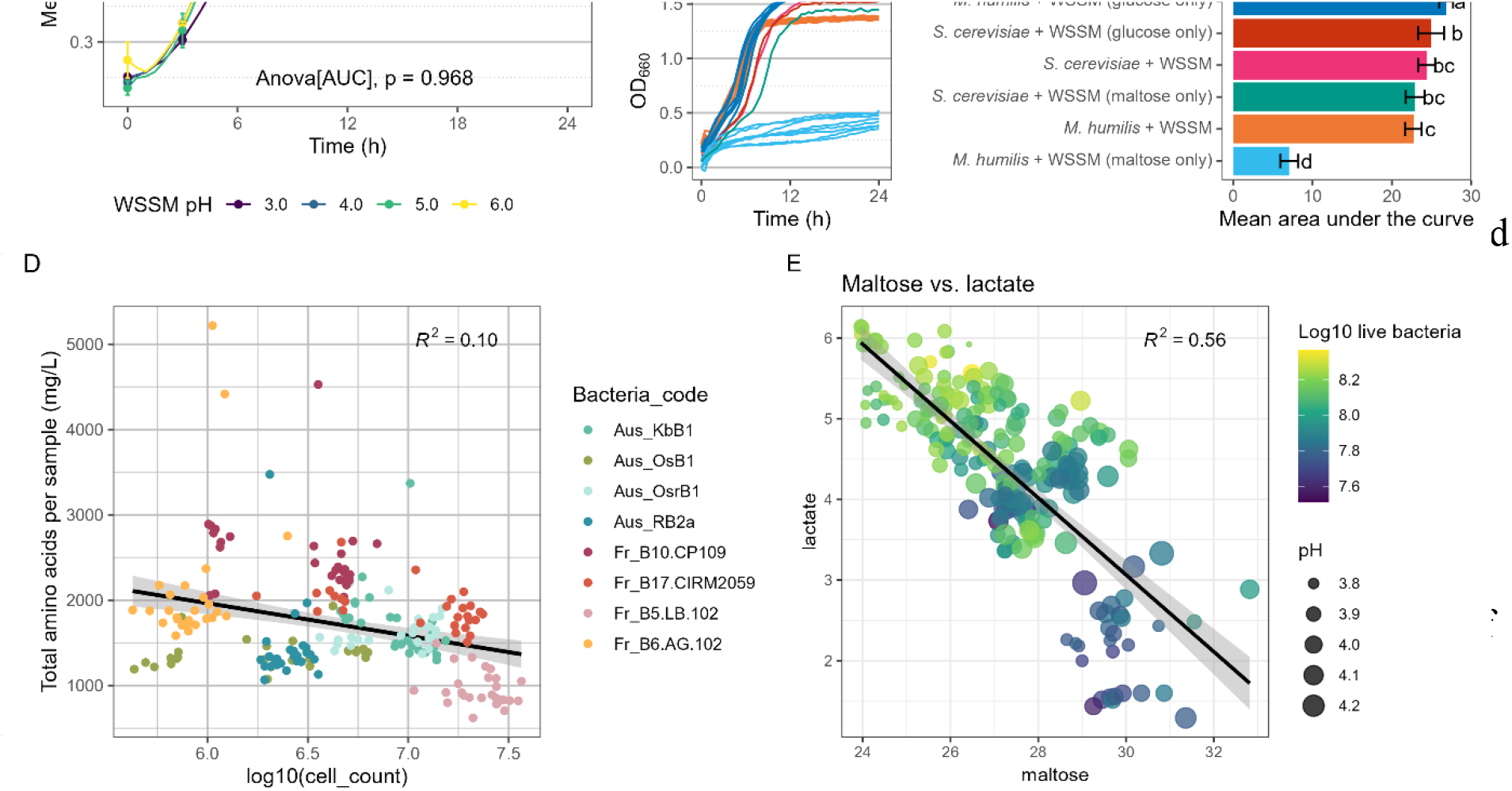
Explaining yeast and bacterial fitness by testing growth in modified media. **A** Modifying WSSM pH to test growth curve response of *M. humilis* Aus_OsrY1. **B** Modifying WSSM carbohydrate composition to test growth curve responses of *M. humilis* strains with reference to *Saccharomyces cerevisiae* Fr_MTF 3622. **C** Searching for a correlation between amino acid accumulation and counts of dead *F. sanfranciscensis* cells. **D** The relationship between maltose, lactate, live *F. sanfranciscensis* counts, and WSSM pH.

## Discussion

The frequent co-occurrence of *M. humilis* and *F. sanfranciscensis* in sourdough starters is often attributed to reciprocal cross-feeding: the bacterium is thought to supply glucose to the maltose-negative yeast (29), while the yeast, in turn, is proposed to provide amino acids to the bacterium (14). However, no studies have clearly demonstrated that glucose produced by *F. sanfranciscensis* from maltose metabolism is taken up by *M. humilis* cells (17). Moreover, the potential of sourdough lactic acid bacteria to provide amino acid to sourdough yeasts has not been investigated. The primary objective of the present study was to test for these two hypotheses by analysing their interaction in a simulated sourdough context. To do this, we assembled a collection of eight yeast-bacteria strain pairs sourced from Australian and French sourdough starters. We compared cell density of strains grown separately and together, and measured concentrations of key metabolites in the spent cell-free medium. The advantage of assessing this interaction in a simulated sourdough rather than a real dough system is the absence of flour amylases that continuously contribute to substrate maltose conditions.

Counts of dead *M. humilis* cells significantly increased when grown with *F. sanfranciscensis*, whereas *F. sanfranciscensis* growth was unaffected by the presence of any co-culture *M. humilis* strain (**Figure 1**). Given their hypothesised complementary dough carbohydrate metabolic properties and their frequent co-occurrence, we expected that the presence of *F. sanfranciscensis* would allow *M. humilis* to benefit from the maltose present in a simulated sourdough, at least to some extent. Sourdough-derived *M. humilis* strains were unable to use maltose for growth, instead demonstrating a clear preference for glucose (**Figure 2**A). *S. cerevisiae*, however (to which *M. humilis* is closely related) is capable of maltose fermentation due to its expression of *MAL* loci, which each encode a regulator and genes required for maltose transport and hydrolysis (8, 30). Despite sourdough lineages of *S. cerevisiae* having increased *MAL* copy numbers relative to non-bakery strains, indicating that selection pressure for maltose assimilation in bakery yeasts exists, we found that the genome of the eight *M. humilis* tested strains does not include *MAL* homologues (31). *M. humilis* therefore does not compete with *F. sanfranciscensis* for this resource (**Figure 2**A). The temperature of the co-culture experiment (25 °C) should have favoured growth of *M. humilis*, as its temperature optimum (27-28 °C) is slightly lower than that of *F. sanfranciscensis* (32-33 °C, (10, 20, 32)). *F. sanfranciscensis* strains could possibly have secreted compounds to kill the yeast cells in a bid to out-compete them, however such mechanisms have not yet been described.

In addition to comparing metabolites in mono- and co-cultures, we also compared concentrations of carbohydrates and amino acids in un-inoculated medium with those in spent media (**Figure 2**). Glucose levels were exhausted in both treatments containing *M. humilis* (**Figure 2**A), but maltose was depleted to a lower extent than predicted in co-culture (**Figure 3**), suggesting that *F. sanfranciscensis* may have accounted for some of the co-culture glucose consumption. This is supported by *F. sanfranciscensis* being able to take up both glucose and maltose (**Figure 2**). In dough, it has previously been found that glycerol levels are significantly increased when yeast strains are present (17). This somewhat aligns with our data from simulated sourdough, which showed greater enrichment of glycerol in co-culture and yeast monoculture than in bacterial monoculture. The production of acetate by *F. sanfranciscensis* in co-culture with *M. humilis* was significantly higher than predicted based on monoculture data (**Figure 3**). This is potentially reflective of a shift in *F. sanfranciscensis* carbohydrate metabolism in response to the presence of *M. humilis*, and may also be linked to bacterial strain identity as well as maltose consumption (17, 18).

Amino acid accumulation in spent WSSM was mainly associated with the presence of *F. sanfranciscensis*, whereas amino acids were largely depleted in *M. humilis* monocultures (**Figure 2**B). Conversely, fermentation by *F. sanfranciscensis* (then referred to as *Lactobacillus brevis* var. *lindneri*) in sourdough led to the accumulation of amino acids, but when co-cultivated with *Candida krusei* or *S. cerevisiae* the total amino acid content was reduced (33). Here, lysine was depleted in all monoculture and co-culture treatments, indicating that both *M. humilis* and *F. sanfranciscensis* were using lysine for protein biosynthesis (34). While the predicted co-culture consumption not significantly differ from its measured values (**Figure 3**), *F. sanfranciscensis* may still have consumed more of the lysine available. Serine and threonine were enriched in bacterial monoculture treatments but depleted in the other two types containing *M. humilis*. Further, threonine depletion in co-culture was greater than predicted (**Figure 3**), indicating an increase in *M. humilis* consumption, or a reduction in *F. sanfranciscensis* production, possibly as a competitive mechanism. Conversely, Ponomarova *et al*. found that serine and threonine were consumed by *Lactiplantibacillus plantarum* when produced by *S. cerevisiae* (15). Similarly to threonine, the glutamine concentration in co-culture treatments was lower than predicted, and while this study found no indication of glutamic acid or glutamine cross-feeding occurring, Ponomarova *et al*. demonstrated that adding these amino acids to a medium in which the LAB was previously unable to grow fully restored their growth (15). This contrast with our findings indicates that if any amino acid-based cross-feeding is occurring between *F. sanfranciscensis* and *M. humilis*, it is likely to differ from previously observed similar phenomena in both the identity of the amino acids involved and the direction of cross-feeding.

In general, co-culture data cluster according to the strain of *F. sanfranciscensis* used (**Figure 4**). This reflects our finding that the degree to which yeast mortality was increased depended on which *F. sanfranciscensis* strain was present in co-culture (**Figure 1**D). Altilia *et al.* also showed that both co-culture growth rate and the ratio of bacterial to *M. humilis* cells were strongly affected by *F. sanfranciscensis* strain (19). These differences are likely to be at least partially a function of each strain’s ability to assimilate and metabolise maltose. In a related study, *F. sanfranciscensis* TMW 1.2138 failed to compete against other strains due to its weak maltose fermentation, as well as the absence of glucose fermentation (18).

Furthermore, the fitness and metabolism of *F. sanfranciscensis* in co-culture was directed predominantly by strain (**Figure 4**). The strain-based variation we observed was reflected in Carbonetto *et al.*, who showed slight differences in CFU per g bread dough after 24 h between two *F. sanfranciscensis* strains (17). The individual capabilities of *F. sanfranciscensis* strains to metabolise maltose and glucose were likely to be linked to strain growth performance, as slight medium composition effects were also observed (data not shown). This aligns with the literature; genome annotation of a collection of 24 *F. sanfranciscensis* strains isolated from sourdoughs in Europe and the USA revealed intraspecific differences in sugar metabolism (maltose, sucrose, and fructose) and the use of electron acceptors (35). Maltose utilisation by *F. sanfranciscensis* is controlled by one or two operons, and variations in their expression and identity may contribute to the strain-based growth differences observed (35). The operon containing *mapB*, which is usually found in sourdough-derived isolates either alone or in conjunction with the *mapA* operon, encodes a major facilitator superfamily (MFS) transporter, a maltose phosphorylase, an epimerase, and a phosphoglucomutase (36). This operon is constitutively transcribed, whereas promotion of the *mapA* operon is induced by maltose (36). Furthermore, this variation in carbohydrate utilisation genes echoes broader variation in the genomes of *F. sanfranciscensis* strains, whose pan genome is accounted for by only 43.14% of its core genome (35, 37).

To investigate whether the decrease in pH associated with acidification caused by *F. sanfranciscensis* was contributing to *M. humilis* mortality, we tested growth of one strain in four pH-adjusted media (**Figure 5**A). Our results show that at least for the strain tested, the pH of WSSM adjusted with lactic acid does not impair yeast fitness. This agrees with previous results, as media-based trials showed that *M. humilis* growth in mMRS (modified for yeast growth by substituting maltose and fructose for glucose) was unaffected by pH values ranging from 3.5 to 7 (20, 32). *F. sanfranciscensis*, however, is affected by pH, and has a clear growth optimum at approx. 5.25 (20). It is thought to use DNA repair enzymes and proteases to mitigate the metabolic stress resulting from its organic acid production (17, 35).

Spent medium pH was associated with the concentration of glucose and particularly maltose remaining in the medium, which were in turn associated with numbers of living *F. sanfranciscensis* cells both in mono- and co-culture (**Figure 5**D). Carbonetto *et al.* have suggested that in a co-culture dough system, maltose may be preferentially converted into organic acids, ethanol, and CO_2_ by LAB rather than glucose for yeast consumption (17). In both our and previously published data, accumulation of these metabolites acted as an indicator of bacterial metabolism occurring (18). We observed no effect of the presence of *M. humilis* on maltose levels in WSSM, as no starch was present in the medium to act as a substrate for amylase (17). The *M. humilis* strains tested did not vary in their inability to grow on maltose, either (**Figure 5**B). Intraspecific genetic variation of *M. humilis* is therefore unlikely to be related to maltose metabolic capabilities (37).

In the synthetic sourdough medium context studied here, we did not find that the presence of *F. sanfranciscensis* allowed *M. humilis* to benefit from the maltose present in the medium. *M. humilis* and *F. sanfranciscensis* have been described as existing in a mutualistic relationship supported by glucose cross-feeding on multiple occasions (35, 38–40). Other studies and reviews have suggested that this relationship is possible because the maltose phosphorylase activity of *F. sanfranciscensis* leads to glucose accumulation in the medium/substrate (13, 29, 41–45). Gobbetti *et al.* co-cultured *M. exigua* (which is maltose-negative) and *F. sanfranciscensis* in MRS supplemented with flour carbohydrates as well as a wheat flour hydrolysate medium and demonstrated that co-culturing the yeast and bacteria affected bacterial cell yield and organic acid production, with a strong focus on differences in maltose metabolism (16). Nonetheless, published evidence that *M. humilis* consumes the glucose produced by *F. sanfranciscensis* in real or simulated contexts is scarce.

A key limitation of this study is that it only covered 24 h of co-cultivation of *F. sanfranciscensis* and *M. humilis*, and refreshments of sourdough starters both for the purpose of maintaining the starter and producing sourdough bread can be longer than this (46). Altilia *et al.* continued their co-culture growth assay past the initial 24 h by passaging cell suspensions into fresh medium a further two times, and found that the refreshment iteration affected growth rate and lag phase (19). Furthermore, growth temperature clearly affects both yeast and bacterial growth, especially as the optima of *M. humilis* and *F. sanfranciscensis* differ, so testing their interaction over a range of temperatures would yield greater insights. We also did not include a treatment that physically separated yeast and bacterial cells in our experiments, and direct cell-cell contact is known to affect LAB gene transcription levels (16, 47).

Using the approaches described here, we show that more factors explain the relationship between *F. sanfranciscensis* and *M. humilis* than glucose cross-feeding, and that the interactions taking place in sourdough starters are likely to be more complex than previously described. The roles of amino acids in this interaction have been examined for the first time, and this examination has revealed that patterns observed between *S. cerevisiae* and other lactic acid bacteria are not always analogous to those between *M. humilis* and *F. sanfranciscensis*. Further work that explores the limitations outlined here, and that more precisely tracks inter-microbial metabolite exchange and underlying genetic and transcription mechanisms would yield more detailed insights into the interaction.

## Conclusions

This study investigated the nature of and possible mechanisms underpinning the interaction between sourdough isolates of *M. humilis* and *F. sanfranciscensis*. This was accomplished using a synthetic sourdough medium which allowed us to quantify both living and dead cells, as well as key carbon and nitrogen compounds linked to potential cross-feeding mechanisms. The data demonstrate that this yeast-bacteria interaction is likely to be neutral, or even competitive, with the strain identity of *F. sanfranciscensis* playing a defining role in the observed dominance of the bacteria and spent medium metabolite composition. The increase in *M. humilis* mortality observed in the presence of *F. sanfranciscensis* may be partially explained by competition for glucose or specific amino acids. This work offers an important point of comparison for previous studies conducted in dough. Future research should focus on the impact of direct cell contact between yeasts and bacteria on their interactions. The present study yields useful observations of a core sourdough interkingdom pairing, which can be exploited to modulate bread-making outcomes. Here we also contribute to the understanding of how microbial consortia may be stabilised through non-collaborative means.

## Acknowledgements

This work was supported by an Australian Government Research Training Program Scholarship provided by the Australian Government and the University of Melbourne to Anna Wittwer.

The authors wish to thank Thibault Nidelet for his help with data visualisation and analysis.

