## Supplementary figures and images for "Lactic acid bacterium *F. sanfranciscensis* impairs fitness of yeast *M. humilis* in a synthetic wheat sourdough medium"

### Fig S1

#### Gating strategy

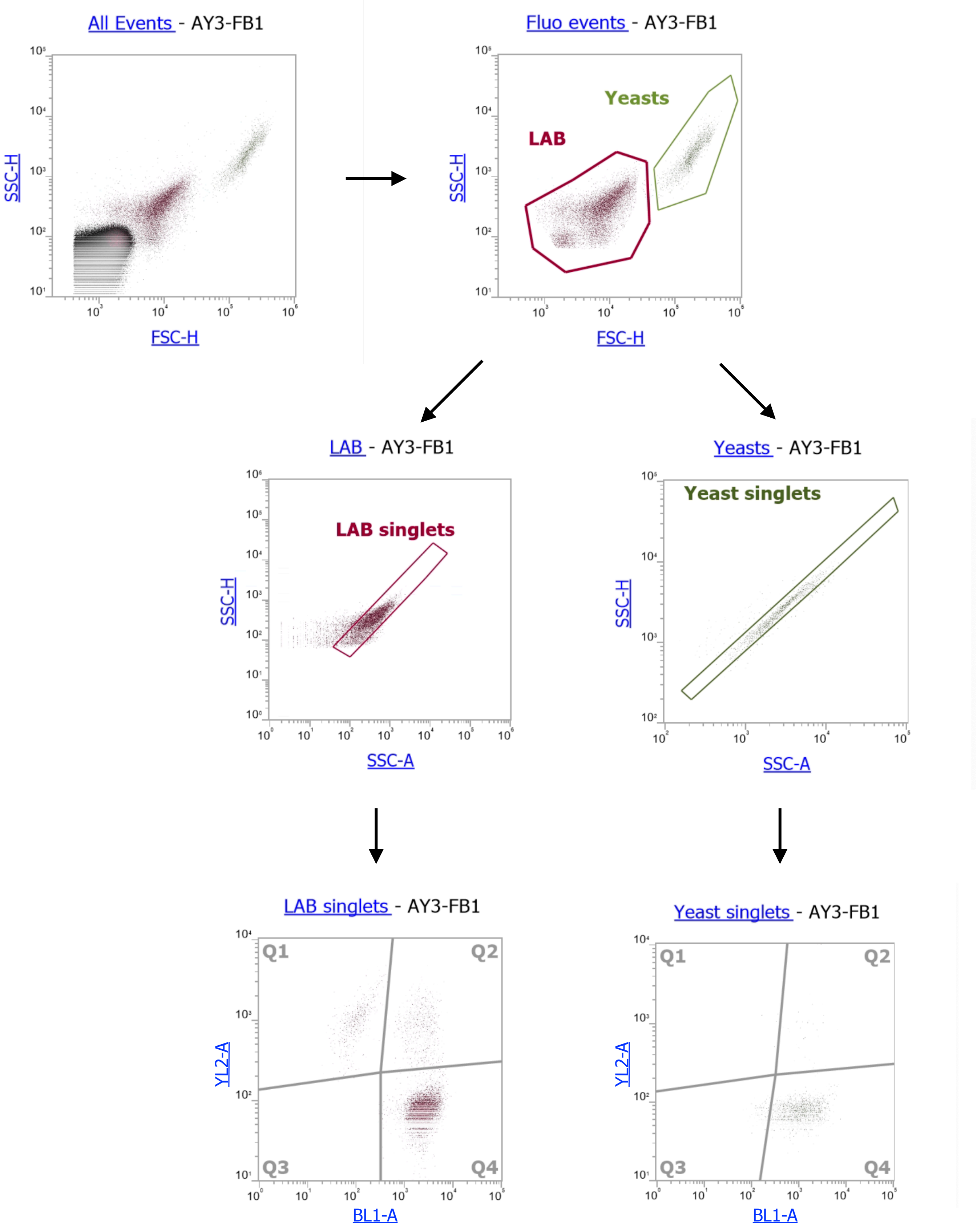

### Fig S2

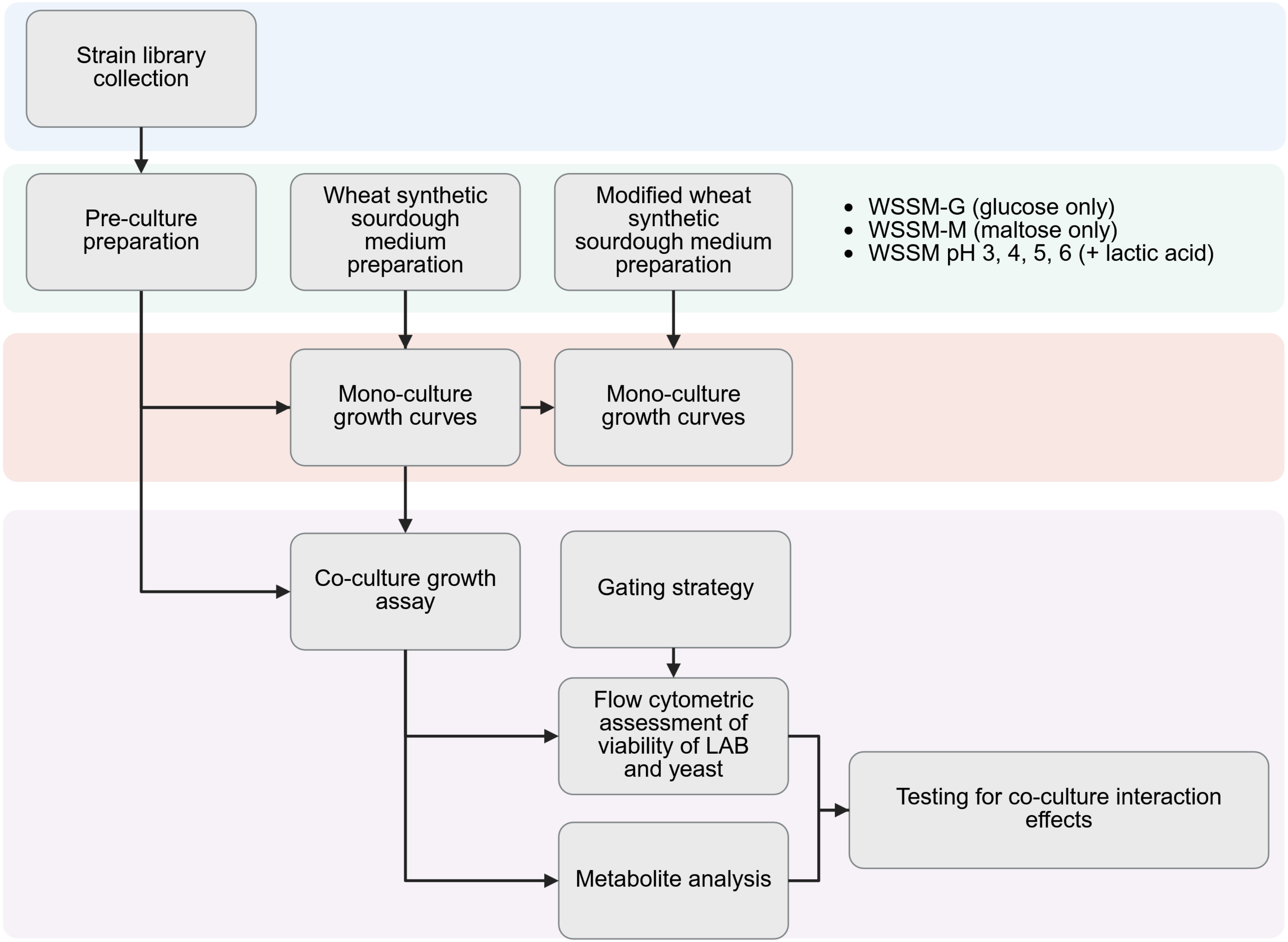
